## Supplementary Figures for "Newly discovered alleles of the tomato antiflorigen gene *SELF PRUNING* provide a range of plant compactness and yield"

**This PDF file includes**

**Figures S1 to S2**

**Legends of Tables S1 to S8**

### Figure S1

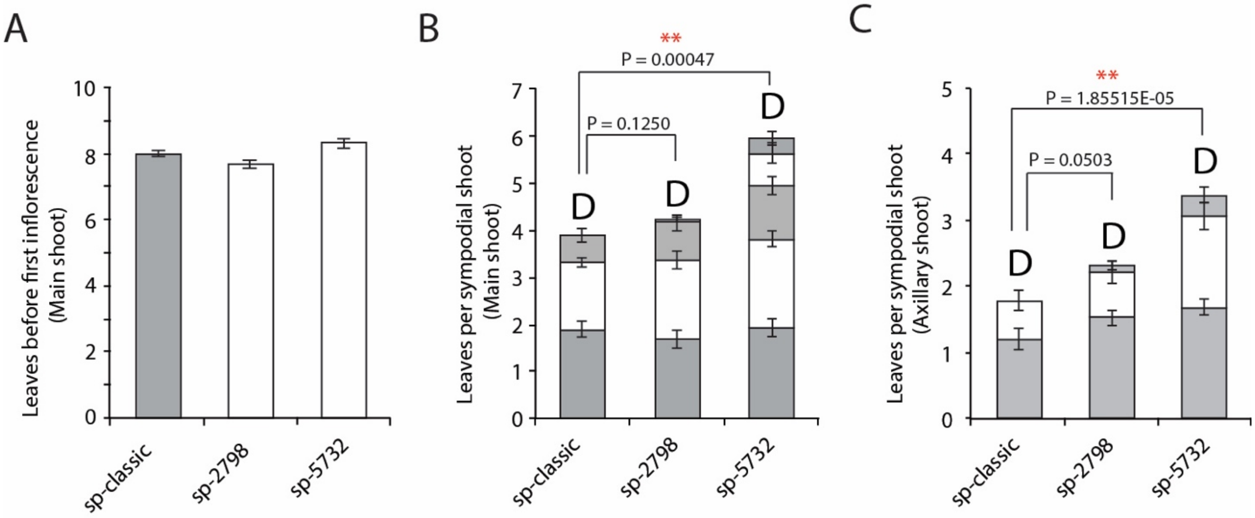


**Supplementary Figure 1.** Comparison of leaf numbers on primary and sympodial shoots among BC3F3 generations of *sp* alleles: (**A-C**). Quantification and comparison of primary-shoot flowering time (**A**) and sympodial-shoots initially produced by the primary shoot (**B**) and axillary shoot (**C**) in *sp* alleles. Statstic analyses were done with at least 16 biological replicates for each genotype. *P* values were determined via two-tailed, two-sample t-test; $^{**}P < 0.01$. CCs of *sp* alleles backcrossed with cv M82 more than four times.

### Figure S2

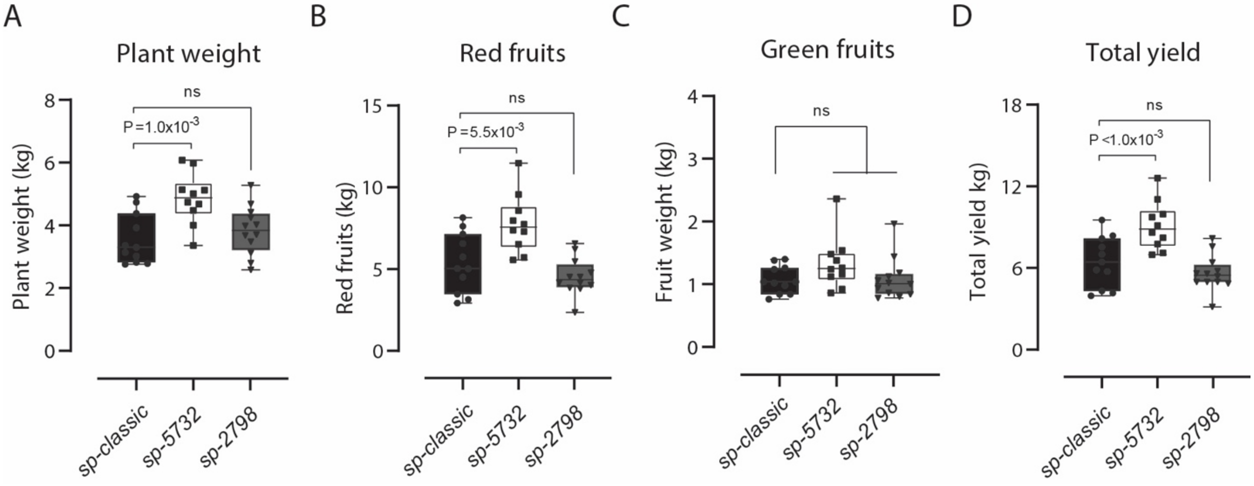


**Supplementary Figure 2.** Quantifications and comparisons of tomato yields among *sp* alleles at the second field trial:

(**A-D**). Statistical comparisons of mean values (±s.e.m.) for plant weight (**A**), red fruit weight (**B**), green fruit weight (**D**), and total yield (**D**) from *sp-classic* as the control (black boxes), *sp-2798* (gray boxes), and *sp-5732* (white boxes). *P* values was determined via two-tailed, two-sample t-test; ns, no significant difference. Statistical comparisons were conducted with more than 10 biological replicates.

**Table S1.** List of Core Collection accessions selected by *sp-classic* genotyping marker

**Table S2.** Genotyping data of *sp* alleles using resequencing data of 588 accessions

**Table S3.** DEGs identified between *SP* and *sp-classic* in TM

**Table S4.** DEGs identified between *SP* and *sp-classic* in SYM

**Table S5.** DEGs identified between *SP* and *sp-classic* in both TM and SYM

**Table S6.** Enriched GO terms of total, TM, SYM, and single-/co-regulated DEGs

**Table S7.** Gene list categorized as developmental process and transcription enriched GO term analysis

**Table S8.** Information of used primers in this study
